# Cryo-FIB lamellae enable ultra-high excitation laser intensities for cryogenic super-resolution fluorescence imaging

**DOI:** 10.64898/2026.04.20.719589

**Authors:** Soheil Mojiri, Joseph M. Dobbs, Jenny Sachweh, Nestor Castillo, Jonas Ries, Julia Mahamid

**Affiliations:** Molecular Systems Biology Unit, European Molecular Biology Laboratory (EMBL), Heidelberg, Germany; Cell Biology and Biophysics Unit, EMBL, Heidelberg, Germany; Collaboration for joint Ph.D. degree between EMBL and Heidelberg University, Faculty of Biosciences, Heidelberg, Germany; University of Vienna, Max Perutz Labs, Campus-Vienna-Biocenter 5, 1090 Vienna, Austria; University of Vienna, Faculty of Physics, Boltzmanngasse 5, 1090 Vienna, Austria

## Abstract

Super-resolution cryogenic correlative light and electron microscopy (SR-cryo-CLEM) integrates molecular specificity and structure on the single-molecule level, but contemporary approaches require low laser powers to preserve specimen vitrification. Here, we perform SR-cryo-CLEM via single molecule localization microscopy (SMLM) on thin biological lamellae and demonstrate that their geometry and minimally-absorbent composition permit excitation laser powers three orders of magnitude higher than standard cryo-EM grids, allowing improved localization precision and imaging speeds.

## Introduction

Cryogenic electron tomography (cryo-ET) enables three-dimensional (3D) visualization of macromolecules at high resolution within their native cellular contexts^1^. However, electron microscopy (EM) data inherently lack specific molecular identity information, and technical limitations currently prevent the direct recognition of smaller proteins and complexes that make up the majority of cellular proteomes^2^. Cryo-CLEM approaches can complement structural data with molecular specificity by providing localization information for fluorescently-tagged molecules of interest^3^. However, the 200-300 nm diffraction limit of light prevents precise localization of individual target molecules. In contrast, single-molecule localization microscopy (SMLM) techniques localize fluorescent emitters with nanometer precision^4–7^ (∼30 nm at cryogenic temperatures using fluorescent proteins^8^).

Yet, there is a fundamental limitation when performing these experiments on cryogenically-preserved biological specimens^9,10^: laser-induced sample heating. Samples must be kept below 138 K to maintain their vitreous near-native states for downstream high-resolution EM analysis, and the powerful lasers typically used in SMLM (up to 500 kW/cm^2^) can cause devitrification, melting, or sublimation. Recent investigations have determined the limits of safe illumination for specimens vitrified on cryo-EM grids, where intensities up to 1 kW/cm^2^ can be used depending on the optical absorbance and thermal conductivity of the grid support material, the illumination wavelength, and the base temperature of the cryostat^11–13^. In practice, contemporary super-resolution cryo-CLEM (SR-cryo-CLEM) applications are limited to laser intensities on the order of ∼0.3 kW/cm^2^ due to heat conduction, optical absorption, and biocompatibility of support films.

Cryo-SMLM with such low laser intensities suffers from prohibitively long measurement times and limited data quality due to high background and reduced single-molecule switching: at the same excitation intensities, the photo-bleaching quantum yield at cryogenic temperatures is at least two orders of magnitude lower than at room temperature^14^. In combination with low laser intensities, this leads to inefficient bleaching of fluorescence background and off-switching of fluorophores. It also leads to longer lifetimes of single-molecule blinking events, resulting in overlapping point spread functions from overly dense emitters, and thus to reduced resolution. Measurement times are lengthened (by several orders of magnitude^3^) to collect enough blinking events, reducing throughput while increasing the probability of image drift or ice contamination on the specimen.

Since laser-induced devitrification thresholds are now understood to depend on the absorptivity and thermal conductivity of the illuminated material^11–13^, we hypothesized that devitrification could be avoided by exclusively illuminating minimally absorbent cellular regions. The absorption coefficient of cellular material is estimated to be around 300 m^-1^, equivalent to ∼0.006% of incident light at 200 nm sample-thickness^15^. This provides an obvious synergy with sample preparation for cryo-ET by focused ion beam (FIB) milling^16^, where thin (∼200 nm) lamellae of suspended biological material are generated. To examine our hypothesis, we modeled experimentally-informed FIB-lamella geometry (Fig. 1a, b, Methods, Tables S1, S2) and used finite-element simulations to study heat transfer within such samples (Methods). The simulated lamella (Fig. 1a, b), fronted by an organ-ometallic platinum (Pt) layer, was illuminated at a central position (Fig. 1a) with a uniform rectangular laser beam (Methods). Our heat transfer simulations identified devitrification thresholds ranging between 500 and 1500 kW/cm^2^, dependent on lamella thickness (Fig. 1c, d, Methods). These non-devitrifying intensities are 500 to 1500-fold higher than when illuminating EM grid materials^11–13^. However, the addition of a thin conductive Pt sputter layer on top of the lamella, which is occasionally performed to reduce charging in cryo-ET, substantially lowers this threshold to 0.5-2 kW/cm^2^ (Fig. 1e).

**Figure 1:**
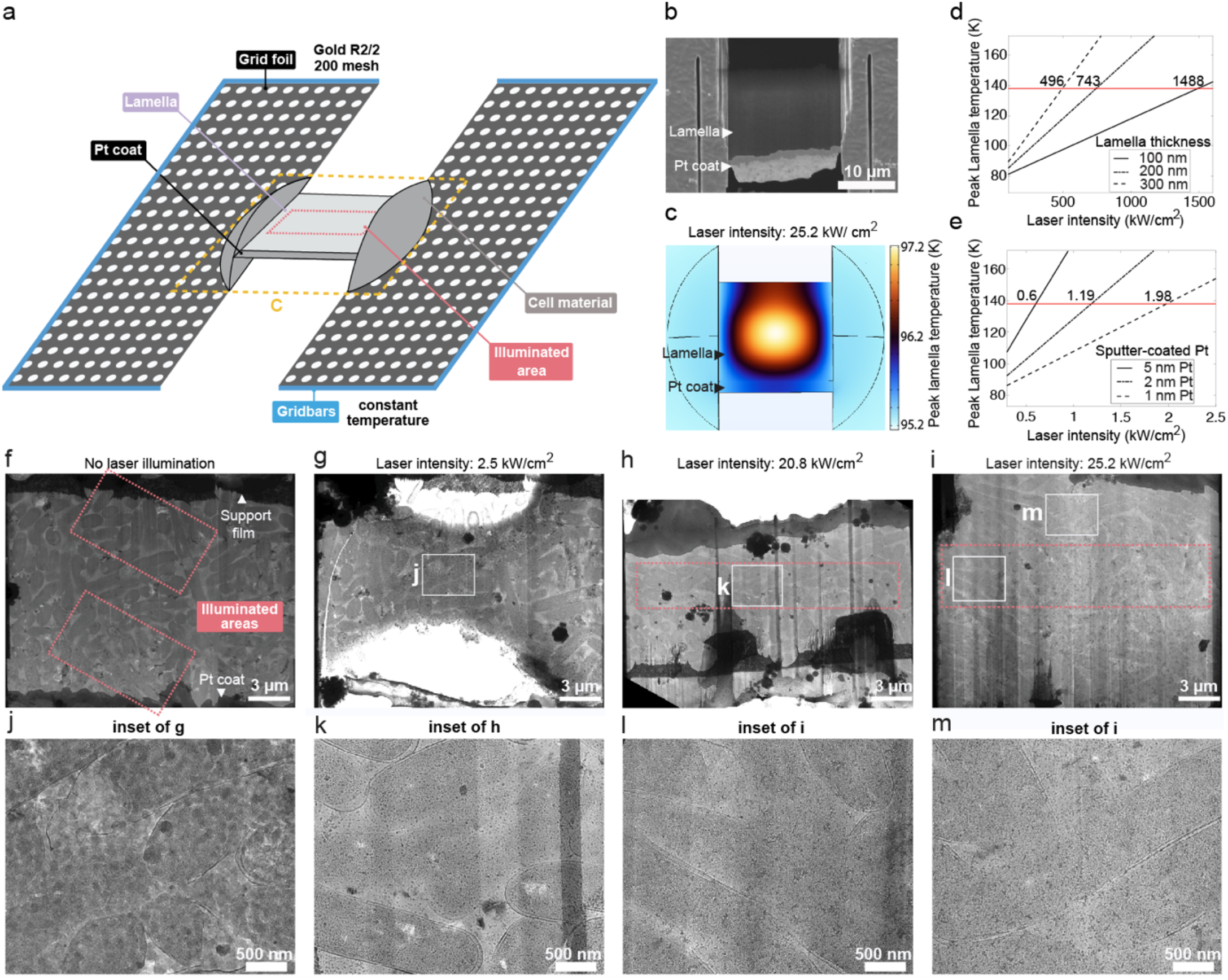
Simulations and experiments demonstrate that site-specific illumination of cellular material in cryo-FIB lamellae enables safe use of high laser powers. **(a)** Simulated geometry of a cryo-FIB lamella at the center of an EM grid square. **(b)** Scanning EM image of a cryo-FIB lamella. **(c)** Simulated temperature profile of the lamella in (a) when illuminated at our maximum experimentally-attainable laser intensity (25.2 kW/cm^2^). Lamella temperature does not approach the devitrification threshold (138 K). **(d, e)** Simulated temperature and devitrification threshold (red line) intensities at 488 nm for lamellae of different thicknesses (d), or for 200 nm-thick lamellae with post-milling conductive Pt sputter layers of different thicknesses (e). **(f-g)** Transmission EM (TEM) lamella maps of *E. coli* cells before (f) and after (g) deliberate illumination of the high absorption support material and protective organometallic Pt. **(h-i)** TEM lamella maps post-illumination at 20.8 (h) and 25.2 (i) kW/cm^2^. No devitrification is apparent. **(j-m)** Magnified images corresponding to the labeled insets in (g-i).

To test our simulations experimentally, we generated FIB-lamellae of *Escherichia coli* cells plunge-frozen on gold EM grids and illuminated these at different laser intensities (Methods, Table S3). First, we deliberately illuminated leftover gold support material at the back of the lamellae, and the protective Pt coats at the front, with a 488 nm laser at 2.5 kW/cm^2^. This resulted in immediate sublimation of the nearby cellular material (Fig. 1f, g). Next, we exclusively illuminated the biological portions of the lamellae. Even at the highest intensity achievable in our microscope (25.2 kW/cm^2^; combination of four lasers, Methods), we neither observed devitrification nor any other laser-induced changes to the sample (Fig. 1h-i, k-m). These experimental data demonstrate that illumination of the minimally-absorbent cellular material increases the safe illumination intensity by at least 25-fold, in line with the 500 to 1500-fold improvement suggested by our simulations.

To demonstrate the feasibility of on-lamella SR-cryo-CLEM with higher laser intensities, we performed a correlated SMLM cryo-ET experiment to visualize the constricting FtsZ Z-ring^17^ in dividing *E. coli*. Lamellae (Fig. 2a) from cells expressing FtsZ-rsEGFP2 (Methods) were illuminated with a 488 nm laser at 2.5 kW/cm^2^ (Fig. 2b, Sup. video 1). We observed localizations at apparent mid-cell constriction sites, but also (potentially nonspecifically) along cell membranes (Fig. 2b). The correlated TEM lamella map (Fig. 2b) contained potential cell division sites (Fig. 2a, insets) that colocalized with distinct patterns of cryo-SMLM FtsZ signal. Indeed, corresponding cryo-ET data (Fig. 2c, Sup. video 2) contained a partial FtsZ Z-ring, which could be visualized and segmented as thin (∼5 nm) filaments traversing the constricting cell-division site (Fig. 2d). The number of localizations (Fig. 2e) and localization precision are provided (Fig. 2f) alongside a comparison to a diffraction-limited rendering of the same region (Fig. 2g, h). Our measurements (Table S4, Methods) indicate that increased excitation intensity reduced the mean fluorophore on time and increased fluoro-phore brightness.

**Figure 2:**
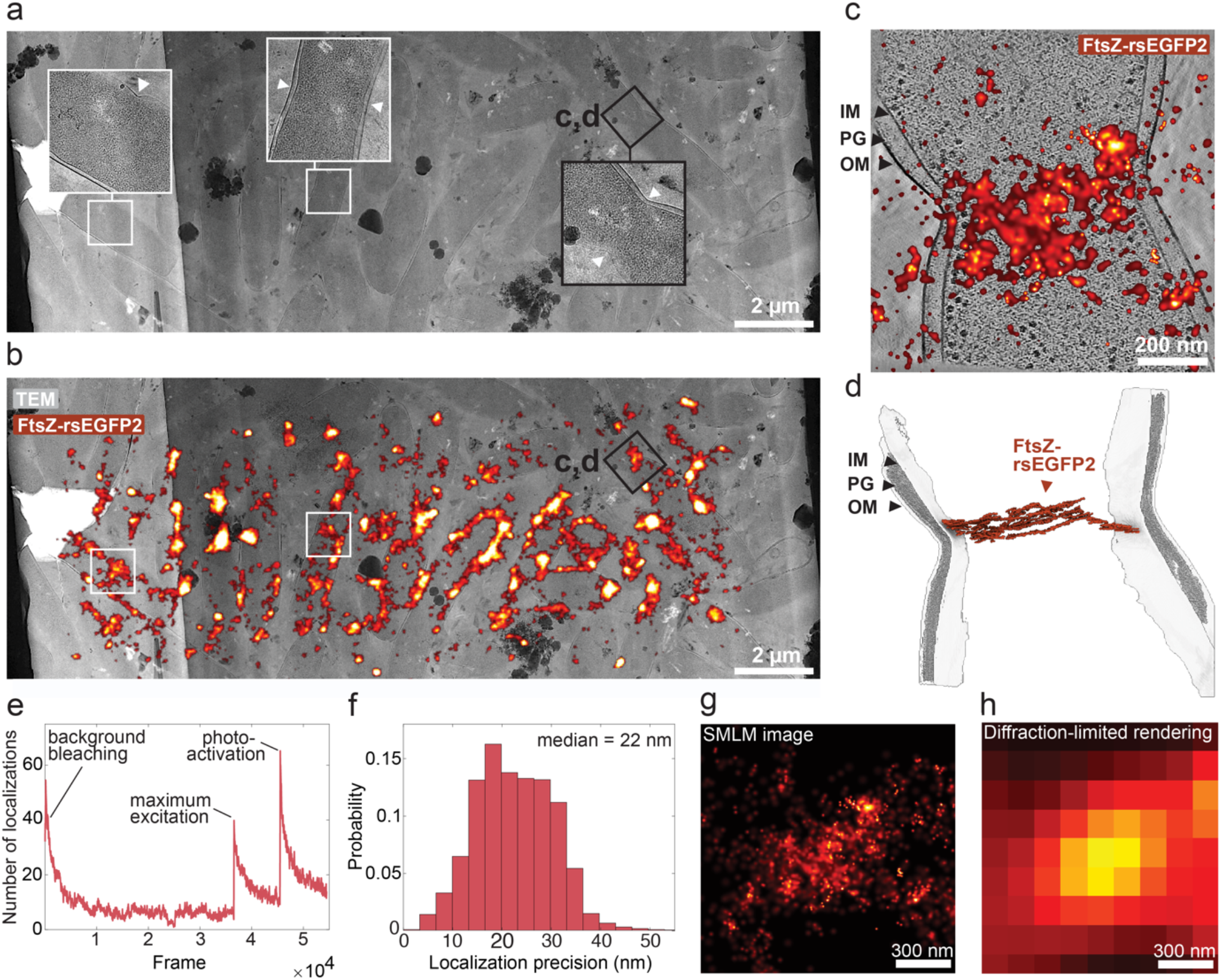
On-lamella SR-cryo-CLEM. **(a-b)** Post-illumination (488 nm, 2.5 kW/cm^2^) TEM lamella map of *E. coli* cells expressing FtsZ-rsEGFP2 (a), overlaid with the correlated SMLM map (b). White arrowheads indicate membrane constriction in magnified insets. **(c)** Slice through a denoised tomographic volume with correlated FtsZ-rsEGFP2 SMLM signal, and **(d)** corresponding segmentation. **(e)** SMLM localizations over time. **(f)** Localization precisions for the entire SMLM map presented in (b). **(g)** SMLM image from (c). **(h)** Diffraction-limited rendering corresponding to (g). IM: inner membrane, PG: peptidoglycan, OM: outer membrane.

In summary, we demonstrate that the unique geometry pro-vided by cryo-FIB lamellae enables the straightforward inte-gration of high-power super-resolution imaging with con-ventional cryo-ET pipelines. By significantly increasing the permissible laser intensity, this approach expands the prac-tical limits of SR-cryo-CLEM.

## Acknowledgements

For technical support and resources, we are grateful to E. Zagoriy, J. Bartho and S. Unger from the EMBL cryo-EM platform, the EMBL Imaging Centre, EMBL IT, and T. Hoffmann. J.M. acknowledges support from the EMBL. J.R. and J.M. acknowledge a Chan Zuckerberg Initiative grant for visual proteomics (2021-234620). This paper was typeset with the bioRxiv word template by @Chrelli: www.github.com/chrelli/bioRxiv-word-template

## Data and code availability

The COMSOL program for simulations (FIB-SEM_Lamella_HeatTransfer.mph) is available at https://github.com/smojiri/LamellaDevitrification-Sim. The script for drift correction in SMAP (driftcorrection_beads.m) is available at https://github.com/jries/SMAP/tree/master/plugins/%2BPro-cess/%2BDrift. The script for image registration (interactive_aligment_tool_Sr_cryo_CLEM.m) is available at https://github.com/smojiri/Correlation-for-Sr-cryo-CLEM-images_Inter-activeManualAlignment. The tomogram, segmentations, and associated SMLM map are available at EMPIAR (EMPIAR-13447), EMDB (EMD-57340), and Bioimage Archive (S-BIAD3196).

## Competing interest statement

The authors declare no competing interests.

## Methods

### Simulation

We performed finite-element heat-transfer simulations in COMSOL Multiphysics (v6.0) to estimate the laser intensities that lead to ice devitrification in cryo-FIB-milled lamellae. Heat transport was described by the steady-state heat equation (eq. 1):

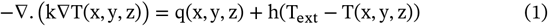

where k is the thermal conductivity, T(x,y,z) is the position-dependent temperature, q(x,y,z) is the volumetric heat source arising from laser absorption, h is the convective heat transfer coefficient, and T_ext_ is the base (cooling) temperature. The laser-induced heat source was modeled (eq. 2) as:

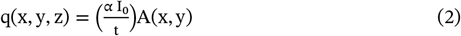

where α is the optical absorption coefficient, I_0_ is the incident laser intensity, t is the lamella thickness, and A(x,y) is the spatial beam profile. To match the field of view of our cryogenic custom SMLM microscope^1^ (laser dosing described below: defined with an adjustable rectangular aperture), the spatial beam profile was represented by a rectangular super-Gaussian (flat-top) distribution (eq. 3), resulting in a near-uniform intensity within the central region and a sharp roll-off at the edges:

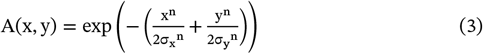

Where n ≫ 2 is the super-Gaussian order, and σ_x_ and σ_y_ define the characteristic beam widths in the x and y directions, respectively. The effective beam widths were defined (eq. 4) using the full width at half maximum (FWHM):

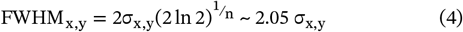

The beam dimensions were set to FWHM_x_ = 18 µm and FWHM_y_ = 7 µm for n ≥ 10, matching the experimentally measured beam size in our custom cryogenic microscope.

The simulated sample geometry represents one square in an R2/2 200-mesh all-gold (*e*.*g*., UltrAuFoil^2^) EM grid, and contains biological material positioned at the center of the grid square. For simplicity, the cellular material here was modeled as a spherical cap with a diameter of 30 µm and a height of 10 µm. A 20 µm × 20 µm × 200 nm FIB-lamella was modelled at the center of this cap with an inclination angle of 12° relative to the grid plane. To represent the typical protective coating layer that is applied ahead of FIB-milling, a 2 µm wide layer of organometallic platinum^3^, with a thickness of 200 nm, was included along the front surface of the lamella. The same layer of 200 nm organometallic platinum was modelled on top of the support film and the peripheral regions of the unmilled biological material. The simulated lamella geometry, while based on a single large cell, is similarly applicable for representations of diverse sample types, including continuous monolayers of small bacterial cells (*e*.*g. E. coli*), clusters of small eukaryotic cells (*e*.*g. Saccharomyces cerevisiae*), and waffle-type^4^ or lift-out lamellae^5,6^ generated from high-pressure frozen specimens.

For the heat transfer simulations, we employed a mesh based on free tetrahedral physics with a fine element size. To quantify the devitrification laser threshold, the primary output of the simulation was the maximum temperure reached within the lamella region. Table S1 describes the thermal conductivity of the materials included in our simulations, and Table S2 describes the optical absorption coefficients of those that were illuminated.

Compared to water, cells absorb visible light more efficiently, which results in stronger heating under laser irradiation^8,9^. To accurately simulate this heating, the linear absorption coefficient of the cells is required. For the simulations, we used an optical absorption coefficient derived from experimentally determined values for MDCK cells (289 ± 114 m^−1^, mean ± standard error of mean [SEM]; n = 7), reported by Huebinger et al.^9^. The reported SEM corresponds to a relative uncertainty of approximately ±39%, reflecting variability across measurements on the same cell. This uncertainty in the absorption coefficient directly results in an equivalent ±39% relative uncertainty in the estimated devitrification laser intensities for lamellae of different thicknesses (Fig. 1d).

We assumed thickness-dependent variations in thermal conductivity to be minimal for vitrified cellular material in the 100-300 nm range (Table S2), and therefore set a constant thermal conductivity for all simulated lamella thicknesses. For lamellae coated with an additional post-milling conductive platinum sputter layer, the optical absorption and thermal conductivity of the vitrified cellular material were negligible relative to the platinum.

The gold support film, which is in thermal contact with the grid bars (the heat sink), was maintained at a constant temperature corresponding to the cryostat base temperature (see below)^10^. The convective heat-transfer coefficient was set to 20 W/mK^2^ to model the presence of cold nitrogen gas surrounding the sample in our open cryostat^10^. For the plots in Fig. 1d and 1e, the base temperature was set to 77 K to represent a generic liquid-nitrogen cryostat. In Fig. 1c, the base temperature was set to 95 K to reproduce the operating temperature of our system. For a lamella at a base temperature of 95 K with a thickness of 200 nm, approximating Fig. 1i, the simulated laser-intensity threshold for devitrification was 545 kW/cm^2^. As room temperature SMLM laser intensities are typically on the order of 500 kW/cm^2^ or below^11^, our simulations indicate that on-lamella SMLM can be performed within similar intensity regimes, and may uniquely benefit from high intensities due to the reduced quantum yield of photobleaching at cryogenic temperatures compared to room temperature.

### *E. coli* preparation and vitrification

The fluorophore rsEGFP2 was selected based on its performance in previous cryogenic SMLM applications^12^. As such, an isopropyl β-d-1-thiogalac-topyranoside (IPTG)-inducible FtsZ-rsEGFP2 plasmid with chloramphenicol selection was constructed using Gibson assembly^13^ via GeneArt master mix (Thermo Fisher Scientific, cat: A46628), from a plasmid expressing FtsZ-mEos2 with a T7 promoter (Addgene #49764, gift from Jie Xiao^14^) and a plasmid expressing rsEGFP2 (Addgene #102879, gift from Stefan Jakobs^15^).

*E. coli* BL21 DE3 cells were transformed with this plasmid and grown to optical density (OD_600_) 0.6 in Lysogeny Broth (LB) medium, at 37°C, with 20 µg/ml chloramphenicol and 300 rotations per minute shaking. IPTG was added at a final concentration of 0.1 mM to induce expression of the fusion construct. After 1 hour, the cells were collected by centrifugation at 5,000 g and resuspended to OD_600_ 32 in M9 minimal medium (for background fluorescence reduction). 3.5 µl of the *E. coli* cell suspension was applied to Ul-trAufoil R2/2 200 mesh gold grids (Quantifoil Micro Tools), which had been plasma cleaned (Fischione M1070) for 1 minute with a 75:25 argon: oxygen gas mix. Grids were vitrified in an ethane/propane cryogen after blotting from the back (1 second, 45% humidity) in a Leica GP1 plunger (Leica Microsystems). Grids were stored in sealed boxes in liquid nitrogen until use.

### Cryogenic FIB-milling and quality control

Lamellae were prepared using an Aquilos FIB-SEM instrument (Thermo Fisher Scientific) with a stage temperature of 83 K. The samples were sputter coated with platinum (30 seconds), coated with organometallic platinum (40 seconds) using the gas-injection system (GIS), and then sputter coated with platinum again (30 seconds), to reduce charging and curtaining. AutoTEM5 (Thermo Fisher Scientific)^16^ was used to prepare lamellae in a stepwise automated fashion, at a 12° milling angle, with a gallium ion beam (currents ranging from 1 nA to 10 pA) at 30 kV.

For cryo-TEM imaging, a Talos TEM (Thermo Fisher Scientific) operated at 200 kV was used to collect lamella maps, using SerialEM^17^, at 3,600x nominal magnification (55.96 Å/px), using a Falcon 3 detector (Thermo Fisher Scientific), a fluence of 0.5 e^-^/Å^2^, a 100 µm objective aperture, and -70 µm defocus. Alternatively, a Titan Krios G3 (Thermo Fisher Scientific) operated at 300 kV was used. It was equipped with a K3 detector and a BioQuantum energy filter (Gatan), and lamella maps were collected at 6,500x nominal magnification (14.02 Å/px), with a fluence of 1 e^-^/Å^2^, a 70 µm objective aperture, and -70 µm defocus. Crystalline ice was sometimes observed in the medium between individual cells, dependent on the thickness of the plunge-frozen *E. coli* layer, and we proceeded with subsequent laser dosing where no apparent excess of crystalline ice was observed.

### Laser dosing

Our simulations indicated that imaging of biological lamellae should permit laser intensities three orders of magnitude higher than those possible for SR-cryo-CLEM applications, where EM grid support material is illuminated (∼0.05 kW/cm^2^ for carbon support films, ∼0.2 kW/cm^2^ for gold supports at 488 nm^10,19^). For experimental laser dosing, pre-screened lamellae were therefore illuminated for 1 minute using different laser intensities. Proper beam positioning was crucial: to precisely define the illumination field of view of our custom-built cryo-SMLM microscope [1], we mounted a rectangular aperture (SP 60, Owis) in a rotating SM1 threaded lens tube (Thorlabs) at a field plane conjugate to the sample plane. This allowed us to resize and rotate the illuminated area to match the length and orientation of each lamella for laser dosing. At the start of each experiment, the beam position and orientation were verified by illuminating the lamellae at low excitation intensity (∼5 W/cm^2^).

Laser dosing experiments were performed on four grids (labeled I-IV in Table S3), each prepared in a separate session. The laser illumination effects were subsequently assessed by cryo-TEM, as described above. In total, eighteen lamellae were analyzed: one from grid I, five from grid II, five from grid III, and seven from grid IV. Across all experiments with practically achievable laser intensities (up to 25.2 kW/cm^2^), laser-induced devitrification or sublimation of lamellae was only observed when the organometallic platinum or leftover support material were illuminated. Illumination of the front-side organometallic platinum layer or adjacent unmilled thick cellular regions at the lamella edges is expected to induce devitrification at or below the threshold reported for carbon-supported grids (< 50 W/cm^2^) [10, 18, 19], due to the strong optical absorption of organometallic (or sputtered) platinum.

We achieved the maximum laser intensity, here 25.2 kW/cm^2^, by combining all our available lasers: 405 nm (2.5 kW/cm^2^), 488 nm (2.5 kW/cm^2^), 561 nm (14 kW/cm^2^), and 640 nm (6.2 kW/cm^2^). For the dosing experiments, lamellae were mounted in our custom-built cryogenic microscope for substantial lengths of time, eventually leading to mild surface ice contamination; the lamellae illuminated at 20.8 kW/cm^2^ (Table S3), were mounted in the cryostat for 9 hours and were relatively clean (Fig. 1h, k), but the lamellae illuminated at 25.2 kW/cm^2^ (Fig. 1i) were mounted in the cryostat for more than 12 hours, and had visible ice contamination (Fig. 1l, m). It may also be possible that ice contamination could have occurred during microscope loading or unloading. For the SR-cryo-SMLM experiment in Fig. 2c, lamellae were mounted in the cryostat for nearly 6 hours. These appeared to have little to no ice contamination.

### Effect of excitation intensity on single-molecule photophysics

To assess the effect of increased excitation intensity on cryo-SMLM performance, we compared single molecules on two independent lamellae containing FtsZ-rsEGFP2-expressing cells. One lamella was imaged with 488 nm excitation at 0.5 kW/cm^2^, and another at 2.5 kW/cm^2^, both substantially above determined devitrification limits for carbon or gold grids^18^. In both cases, an initial conditioning step was carried out, consisting of 1 hour of illumination at 0.25 kW/cm^2^ (488 nm) to drive fluorophores into the dark state and bleach the autofluorescence background. The field of view and camera exposure time (100 ms) were kept constant to ensure direct comparability between conditions, and data were acquired through a 515/30 nm emission filter. Single-molecule parameters were quantified over 9,000 frames (Table S4).

### Cryogenic SMLM

The *E. coli* lamellae containing cells expressing FtsZ-rsEGFP2 were prescreened to ensure proper vitrification, and imaged using our custom-built cryo-SMLM microscope. At cryogenic temperatures, as previously described for rsEGFP^12^, the transition of fluorophores to the dark/off state is significantly slower compared to room temperature SMLM. This results in a high initial population of active fluorophores^12,19^. Therefore, to achieve effective single-molecule localization, *i*.*e*. a sparse distribution of emitters per frame, the lamellae were pre-illuminated with a 488 nm laser at 250 W/cm^2^ for 60 minutes to drive most fluorophores into the “off” state. Additionally, prolonged low-intensity illumination helps reduce autofluorescence background by gradual photobleaching. The resulting photobleaching trend is visible in the first 36,000 frames of Fig. 2e. Subsequently, rsEGFP2 was excited at the maximum available 488 nm intensity (2.5 kW/cm^2^), and 18,000 frames with 100 ms exposure time were acquired through a 515/30 nm emission filter. As a result of the increase in excitation laser intensity (after bleaching finished, frame 36,000), a clear rise in the number of detected localizations could be observed (Fig. 2e). The final 9,000 frames were acquired under continuous photoactivation with a 405 nm laser at 5 W/cm^2^. This illumination converts fluorophores from the dark/off state into the active emitting state, generating new single-molecule events and thereby sustaining a sufficient number of localizations throughout the imaging sequence.

Lateral drift correction, single-molecule localization, and downstream analysis were carried out using the SMAP plugin in MATLAB^20^. Raw SMLM movies (TIFF) were imported, and the effective pixel size was set to 136 nm. Single-molecule localizations were detected using the SMAP default peak finder, and fitted using the 2D Gaussian fitter with default fitting and filtering settings, unless stated otherwise. Localization precision was estimated from photon counts and background noise in SMAP.

Sample drift was corrected based on the fiducial markers (beads) algorithm in SMAP^20^ (‘*Drift correction Beads’* plugin), where the x/y drift trajectory over time was estimated, smoothed using a sliding-window mean filter, and subtracted from all localizations. A region of interest containing a single bright point-like feature emitting throughout the rendered data was selected. After drift correction, fluorophores in consecutive frames were grouped (merged) if they were closer than 35 nm and dark for maximally 1 frame.

Localizations were rendered in SMAP^20^ using a red-hot lookup table with a localization precision filter of 0-30 nm and a PSF size of 350 nm. Poorly fitted localizations were rejected using a filter on the normalized log-likelihood. For correlated SR-cryo-CLEM (Fig. 2b, c), only localizations acquired after the initial 36,000 frames (after the bleaching phase) were included in the analysis. This was necessary to avoid spurious localizations from the background fluorescence. The rendered SMLM images were exported as TIFF files for correlation with the low-magnification TEM images (see below). The tomogram-correlated SMLM image (Fig. 2c) was exported as a TIFF with pixel sampling matched to the high-resolution cryo-ET slice by adjusting the pixel size of the reconstructed image.

### Correlated cryo-ET data acquisition and processing

A Titan Krios G4 TEM (Thermo Fisher Scientific) operated at 300 kV was used to collect lamella maps using SerialEM^17^, with a post-column Selectris energy filter and Falcon 4i detector (Thermo Fisher Scientific), at 3,600x nominal magnification (34.60 Å/px), a fluence of 0.7 e^-^/Å^2^, a 100 µm objective aperture, and -70 µm defocus. Tilt series were collected starting at 12° pretilt angle to compensate for the lamella milling angle, and a nominal magnification of 53,000x (2.39 Å/px), using a 100 µm objective aperture. Tilts (3.12 e^-^/Å^2^ each) were collected from -48° to +57° with 3° increments using the dose symmetric scheme^21^, and a defocus range of -3 to -6 µm. The total fluence was 112 e^-^/Å^2^.

Tilt series were preprocessed in WarpTools^22^, aligned with AreTomo2^23^, and reconstructed in WarpTools at 15 Å/px. Tomograms were denoised with IsoNet2^24^, and the Z-ring filaments and peptidoglycan layer were segmented in Dragonfly (Comet Technologies Canada Inc.) using a 2.5D 5-slice U-net, then manually corrected. Inner and outer cell membranes were segmented with MemBrain-seg^25^, and segmentations were displayed using ChimeraX v1.8^26^ and the ArtiaX^27^ plugin.

### Image registration

Registration of cryo-TEM data with SMLM maps is an important challenge and currently introduces substantial localization error; further development is required to obtain reliable solutions. Since our study was focused on examining the feasibility of use for high laser doses, we made use of simple custom MATLAB scripts to manually register images based on image features, including cell membranes, cell sizes, and orientations. Specifically, the script was used to register the lamella map (34.60 Å/px), the position of an acquired tomogram (2.39 Å/px), the SMLM localization map (136 nm/px), and a zoomed region of interest from the SMLM localization map. The script applied precise shifts, rotations, flips, scaling, and transparency adjustments.

To adjust background transparency for overlays of the EM and SMLM data (Fig. 2c), the rendered SMLM images were afterwards converted to transparent overlays by generating an alpha mask via RGB intensity thresholding, where near-black pixels were assigned α=0, and all remaining pixels α=1; mask edges were smoothed using Gaussian filtering.

## Supplemental tables

**Table S1.** Thermal conductivity properties of materials used in simulations.

| Material | Thin film thermal conductivity<br>at 138 K (W/m.K) | References |
| --- | --- | --- |
| Organometallic platinum | 7.2 | 3 |
| Vitreous cellular material | 1 | 28,29 |
| Gold support film | 72 | 30,31 |
| Platinum sputter coat | 51 | 30,31 |

**Table S2.** Optical absorption properties of illuminated materials used in simulations.

| Material |  |  |  |  |
| --- | --- | --- | --- | --- |
| Vitreous cellular material (lamellae)<sup>9</sup> | Thickness | 100 nm | 200 nm | 300 nm |
|  | Optical absorption (%) | 0.003 | 0.006 | 0.009 |
| Platinum at 488 nm<sup>32</sup> | Thickness | 1 nm | 2 nm | 5 nm |
|  | Optical absorption (%) | 3 | 5.7 | 12.2 |

**Table S3.** Laser intensities applied to each lamella, and resulting lamella state.

| Laser intensity (kW/cm<sup>2</sup>) | Number of lamellae that remained vitrified | Number of devitrified/sublimated lamellae |
| --- | --- | --- |
| 1.8 | 1 (grid I) | - |
| 2.5 | 1 (grid II), 3 (grid IV) | 1 (support/Pt illuminated), (grid II) |
| 5.7 | 1 (grid III) | 1 (support/Pt illuminated), (grid II) |
| 8.2 | - | 2 (support/Pt illuminated), (grid II) |
| 10.1 | 1 (grid III) | 1 (support/Pt illuminated), (grid III) |
| 20.8 | 1 (grid III) | 1 (support/Pt illuminated), (grid III) |
| 25.2 | 4 (grid IV) | - |

**Table S4.**
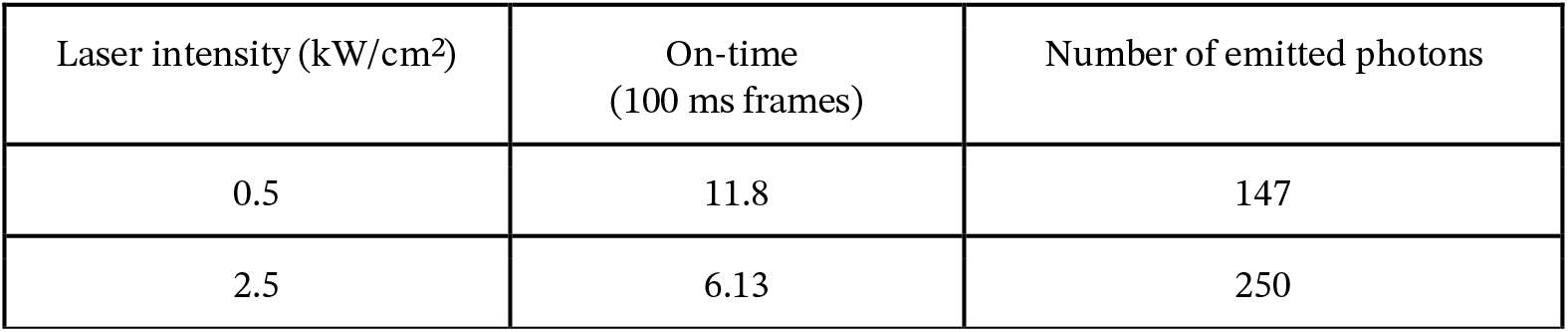
Mean on-time and brightness of single-molecules on lamellae illuminated at different laser intensities. A fivefold increase in excitation intensity reduced the mean fluorophore on-time by approximately twofold while increasing the mean single-molecule brightness by ∼65%.

| Laser intensity (kW/cm<sup>2</sup>) | On-time (100 ms frames) | Number of emitted photons |
| --- | --- | --- |
| 0.5 | 11.8 | 147 |
| 2.5 | 6.13 | 250 |

## Notes

### Competing Interest Statement

The authors have declared no competing interest.

